# An Integrated Method for Profiling Lipid–Protein Interactions Using Multifunctional Lipid Probes

**DOI:** 10.64898/2026.01.17.700062

**Authors:** Scotland E. Farley, Gaelen Guzman, Berit Blume, Frank Stein, Carsten Schultz, Fikadu G. Tafesse

**Affiliations:** Department of Chemistry and Biochemistry, Bates College, Lewiston, ME, USA; Department of Molecular Microbiology and Immunology, Oregon Health & Science University, Portland, OR, USA; Department of Chemical Physiology and Biochemistry, Oregon Health & Science University, Portland, OR, USA; European Molecular biology Laboratory, Proteomics Core Facility, 69117, Heidelberg, Germany

## Abstract

Cellular lipids shape health and disease through specific protein interactions, yet lipid-protein networks remain poorly defined. Despite rapid advances in functional lipid probes, the field still lacks a practical, dedicated protocol for conducting lipid-protein interaction studies. We describe detailed methods for determining lipid interactomes from cells using multifunctionalized lipid derivatives. We provide protocols that detail 1) how to treat cells with lipid derivatives and perform photochemistry to obtain lipid-protein conjugates; 2) how to perform click chemistry with a fluorophore and observe lipid-protein conjugates by in-gel fluorescence; 3) how to perform click chemistry with azide beads and prepare lipid-protein conjugates for proteomic analysis. We provide context on important parameters for each step and include guidelines for controls, as well as suggestions for troubleshooting based on common problems encountered during the preparation of this protocol. This protocol enables mapping lipid interactomes across diverse biological systems. The entire workflow from cell treatment to complete proteomic sample preparation requires ∼15 hours over four days, depending on the type of experimental readout (in-gel fluorescence or proteomics), and the usage of pause points. Practitioners are expected to be familiar with standard biochemical techniques, such as sterile sample handling and tissue culture and gel electrophoresis. Additional skills are needed for mass spectrometric analysis, and collaboration with a proteomics core facility is recommended. The described procedures uniquely enable the identification of the protein interactors (the interactome) of select lipid species, providing for a major shift in the characterization of the biological roles of lipids in cellular systems.

## Introduction

Lipids play multifaceted roles in cellular physiology, serving not only as essential structural components of biological membranes but also as dynamic mediators of signaling pathways. Despite their central roles in cellular physiology, lipid–protein interactions remain less well characterized than protein–protein interactions, due in part to the physicochemical complexity of lipids and the historical lack of compatible experimental tools. The development of synthetic lipid analogs – functionalized lipid probes – has enabled the application of affinity proteomics to lipid interactome studies in biologically relevant systems^1–5^.

Interactions among lipids and between lipids and proteins define key membrane properties such as fluidity, curvature, and detergent resistance, which in turn distinguish the biochemical identities of organelles and membrane microdomains^6^. Beyond structural roles, lipids also function as signaling molecules. For example, sphingosine-1-phosphate (S1P) regulates T cell chemotaxis, while phosphoinositides such as PI(3,4,5)P_3_ control cell survival and growth via effector recruitment^7–10^. These interactions are often transient and spatially restricted, necessitating tools with precise spatiotemporal control.

While gene editing technology has greatly simplified the identification of protein-protein interactions (e.g. via genetic fusion to an affinity handle such as the HA-tag), lipids are not directly genetically encoded and are thus resistant to traditional affinity techniques^3^. Their small size makes them highly sensitive to structural modification, and their rapid metabolic turnover poses further challenges for studying lipid-specific effects^1,6^. Low abundance can also present a challenge: many bioactive lipids such as sphingosine, ceramide, and phosphatidic acid are maintained at very low intracellular concentrations and have half-lives of only minutes^2,11^.

To address these challenges, functionalized lipid probes have been developed to mimic native lipid function while enabling biochemical capture of interaction partners. These bifunctional or trifunctional probes typically include: (i) a diazirine group for UV-induced covalent crosslinking (Fig 1, green); (ii) a terminal alkyne for click chemistry–based enrichment (Fig 1, blue); and (iii) in trifunctional variants, a photolabile cage for temporal control over lipid activity (Fig 1, orange). This design allows for selective stabilization and enrichment of lipid-binding proteins with minimal perturbation to native lipid function. In addition, some lipids that carry extremely dense negative charge, such as phosphatidylinositol phosphates, require masking acetoxymethyl groups to allow them to passively diffuse across the plasma membrane (Fig 1, purple).

**Figure 1:**
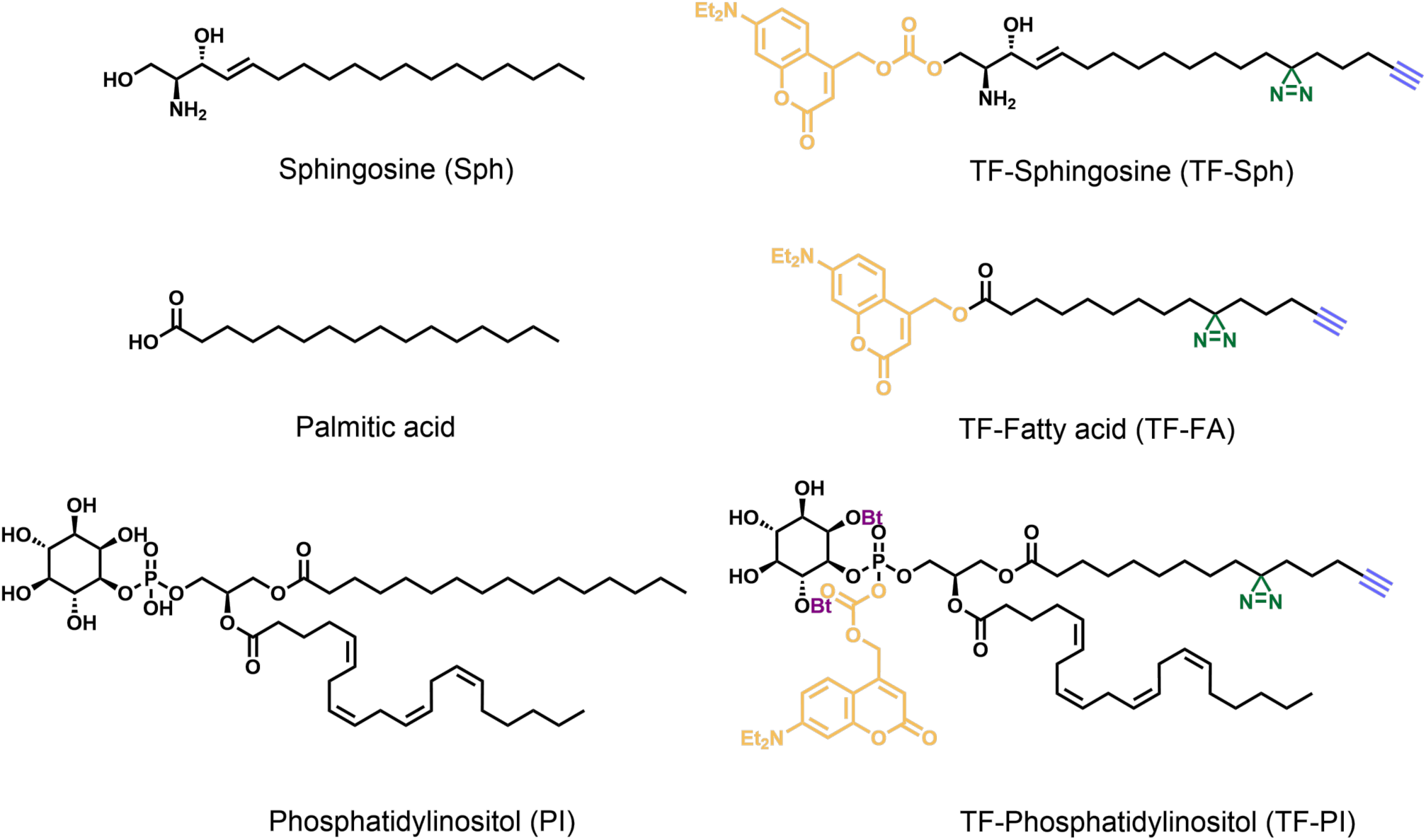
Chemical structures of representative lipid classes: sphingolipids, fatty acids, and phosphatidylinositol’s. The native structures are shown on the left and the trifunctional derivatives shown on the right. The Butyryl ester (Bt, purple) serve as protection groups for the negatively charged lipids to support cell entry and avoid aggregates. The coumarin photocage (orange) protects the lipids from premature metabolism. Diazirines (green) form reactive intermediates upon ultra-violet light irradiation to covalently bind nearby proteins. Terminal alkynes (blue) are used for click chemistry to either conjugate dyes or affinity tags.

This protocol provides a detailed guide to the use of bi- and trifunctional lipid probes in interactome analysis. We review the design and application of each functional group, considerations for experimental design, recommended quality control measures, and strategies for data analysis and validation. We also present a complete protocol for conducting lipid-based affinity proteomics from probe treatment and photoactivation to crosslinking, enrichment, and mass spectrometry sample preparation. As no significant alternative methodologies exist to identify the protein interactors of select lipid species, this protocol represents a major step forward in elucidating lipid interactomes and characterizing the biological activity of select lipids.

### Functionalized Lipid Probes: Photochemistry and Reactivity

#### Diazirine Crosslinking for Capturing Lipid–Protein Interactions

Lipid–protein interactions are often transient and low-affinity, owing to the small size and limited hydrogen bonding capacity of lipids^12,13^. Covalent crosslinking transforms these fleeting interactions into stable adducts suitable for enrichment and mass spectrometry. Among photoactivatable groups, diazirines are favored for lipid probes due to their minimal steric footprint and broad reactivity profile^14^. Upon UV (∼355 nm) exposure, diazirines form reactive carbenes capable of inserting into nearby C–H, N–H, O–H, S–H, or π-bonds. While generally nonspecific, there can be some preference for glutamate residues for diazirines where the diazo intermediates are preferred over the carbene^15^ (Fig 2A).

**Figure 2:**
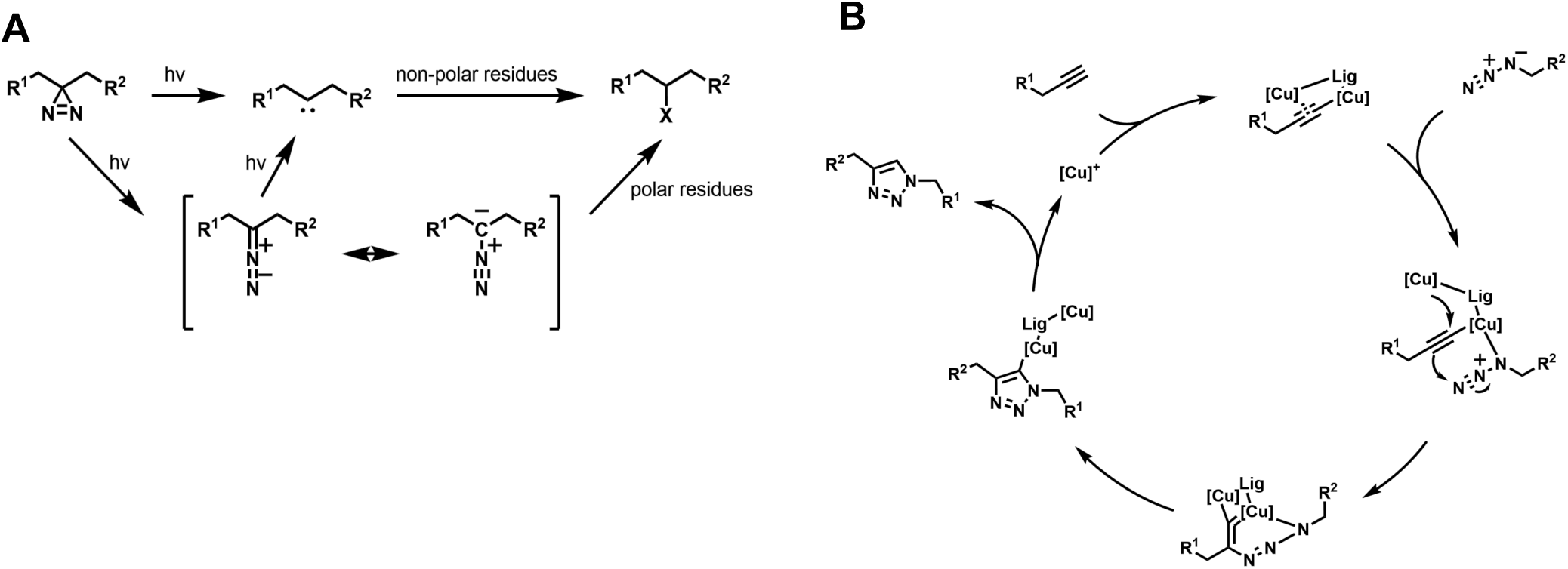
Photoaffinity labeling and click chemistry. A) Photoaffinity labeling is a key method to identify biomolecular interactions and small molecule targets in a cell^1^. Upon irradiation of diazirines diazo and carbene intermediates are generated in sequential order. Whereas the diazo intermediates preferentially react with polar residues, the carbene intermediates react with non-polar residues. This selectivity can be enhanced though power density and irradiation time, a prolonged irradiation leads to increased formation of carbene and therefore a higher selectivity for non-polar residues^2^. B) The copper-catalyzed 1,3-cycloaddition, now known as the click reaction, was published by Rostovtsev et al. in 2002 and offers an orthogonal conjugation method compared to the classic amide coupling^3^. The catalyst is a Cu(I) species that can be obtained either directly from Cu(I) salts or by in situ reduction of Cu(II) salts. The alkyne coordinates the copper and forms the first intermediate. One of the copper atoms can coordinate a nitrogen of the azide to form a complex. The other copper atom attacks the terminal carbon of the double bond, resulting in the ring closure of the metal cycle. A reductive ring closure of the triazole and subsequent protonation lead to the triazole and the release of the Cu(I) catalyst^4^. Lig: Ligand.

Importantly, diazirine placement influences crosslinking specificity. Terminal acyl chain positioning (e.g., C10 in 8-3 Fatty Acid) favors interactions with transmembrane proteins, whereas positioning near the headgroup (e.g., C3 in 1-10 Fatty Acid) captures a broader range, including peripheral membrane proteins^16^. Expanding this design space offers opportunities to interrogate both intramembrane and membrane-interface interactomes.

#### Terminal Alkyne as a Bioorthogonal Affinity Handle

Due to the nature of their synthesis and their structural sensitivity, lipids cannot be genetically tagged. The terminal alkyne provides a compact, bioorthogonal alternative that preserves lipid function while enabling downstream modification^1^. It serves as the reactive partner in copper-catalyzed azide–alkyne cycloaddition (CuAAC), a highly specific click chemistry reaction^17^. As copper(I) is cytotoxic, labeling is performed in fixed or lysed samples (Figure 2B).

Alkyne-modified lipids can be coupled to azide-conjugated fluorophores for imaging or to affinity tags for protein enrichment^12,13^. Click-based capture via azide beads or biotin–streptavidin strategies enables selective isolation of crosslinked lipid–protein complexes for proteomic analysis.

#### Photocaged Lipids for Spatiotemporal Control

The rapid metabolism and turnover of bioactive lipids, often within minutes, complicate functional studies in live cells [^18,19^. Photocaged lipid analogs address this by rendering the lipid inert until photoactivation. Upon exposure to light of an appropriate wavelength, the photolabile group is removed, restoring biological activity with high spatial and temporal precision. Photocaged lipids have proven useful for dissecting localized lipid signaling. Höglinger et al. (2015) used a caged sphingosine analog to trigger lysosomal calcium release upon photoactivation, uncovering a link between sphingosine metabolism and intracellular calcium dynamics ^11^.

Caged lipids are stable in cells for at least an hour, which is in stark contrast to the rapid (∼5 minutes) metabolism of the same lipids once they are uncaged^2^.

Among photocages, 7-diethylamino-4-methylene coumarin is well suited for lipid studies. In addition to efficient uncaging via radical cleavage, it offers intrinsic fluorescence for tracking probe uptake^1^. Alternative cages include nitrobenzyl (shorter wavelengths, non-fluorescent) and BODIPY (bulkier but longer-wavelength activation)^3,20^. The coumarin cage is removed when the molecule is exposed to blue (400 nm) light, which is a critical consideration for probes that also contain a diazirine group, which are activated at 350 - 360 nm light, as these two photochemical reactions must happen orthogonally. Chromatic orthogonality can be achieved using narrow bandpass filters; Höglinger et al (2017) demonstrated that the coumarin photocage and diazirine group can be activated independently under these conditions. Chromatic orthogonality can also be achieved by using light sources with discrete wavelengths of emission, as described by ^16^, and below in this protocol (the “Uncaging Lamp” and “Photocrosslinking Lamp” described in the Equipment).

## Limitations

While bi- and trifunctionalized lipid probes are powerful tools for mapping lipid–protein interactions, several limitations must be considered. Any chemical modification, even minimal, alters the lipid’s biophysical properties, potentially affecting subcellular localization, membrane partitioning, and interaction specificity^21–23^. For instance, coumarin-based photocages prevent premature metabolism but also impair normal trafficking; upon uncaging, the probe becomes globally available, enabling non-physiological interactions^24^.

It should be noted that high-intensity blue and UV light are known stressors for cultured cells and thus may introduce cytotoxic effects which perturb the protein-lipid interactions of an un-stressed cell^25,26^. However, the timing of irradiation with blue and UV light described in the procedures below is brief, and cells will be directly harvested following UV-induced crosslinking – each limiting the extent of these cytotoxic effects. Light sources of lower intensity, such as the nail lamps described previously^16^, are another way to minimize cytotoxicity to cells. It is unavoidable, however, that if the experiment of interest is closely associated with the types of cell stress associated with blue light (for example, ferroptosis) and UV light (for example, DNA damage), that the use of such light sources may inherently perturb the system.

Photoactivatable crosslinkers such as diazirines, though minimally disruptive, also introduce bias. Even slight changes in their positioning can shift the interaction profile, as shown by Farley et al. (2024)^16^. Rigorous experimental design is essential. Negative controls (e.g., non-irradiated samples) must be included for each biological comparison and sufficiently replicated-though this often limits multiplexing capacity in quantitative proteomics methodologies such as TMT, iTRAQ, or SILAC workflows.

Additionally, common normalization approaches may exaggerate protein abundance in -UV controls, producing symmetric volcano plots and misclassifying relevant interactors. Statistical filtering is needed to distinguish true targets from artifacts. It is also of course easiest to see the highest-abundance lipid-binding proteins; identifying low-abundance lipid-binding proteins may require some combination of higher amounts of sample and enrichment of relevant cellular fractions (such as membranes or specific organelles).

Finally, these probes have so far only been validated in cultured cell lines. In vivo applications in tissue samples or vertebrates remain limited due to likely high cost and uncertain pharmacokinetics. Small invertebrate models such as C. elegans may be suitable, but probe uptake and labeling efficiency remain untested.

## Experimental Design and Quality Control

We present below three protocols for 1) producing and harvesting lipid-protein conjugates; 2) visualizing protein-lipid conjugates by in-gel fluorescence; 3) preparing samples for proteomic analysis. The details of the experimental design can vary widely, depending on the biological question of interest. Our protocols remain within the realm of mammalian cell culture (rather than bacterial culture, tissues, or organisms); this is not to say that they could not be adapted to other systems, but further optimization would undoubtedly be required. We have furthermore found that every cell culture system (cell line, lipid of interest) requires its own tweaking to discover the optimal amount of probe and density of cells required to produce robust signal. The procedures described below should be adjusted for the particularities of the analyzed cell type (e.g. use appropriate cell culturing plates and media, adjust cell count for particularly large or small cells, adjust timing of plating). Below, we present baseline aspects of quality control that must be taken into consideration before using lipid probes in any specific system.

### Probe Treatment

Effective labeling requires sufficient probe intercalation into the target cells or tissue, which depends on both the type of cells used in the procedure, and the nature of the lipid probe. Lipid probes can be sorted into three rough entry types: **Type 1** lipid probes are relatively water-soluble and readily cross the plasma membrane (aqueous solutions up to 50 µM can be made at room temperature with no detectible precipitation); **Type 2** lipid probes are quite water-insoluble, but can cross the plasma membrane with the help of the mild detergent Pluronic; **Type 3** lipid probes require a masking group on negatively charged head group moieties to ensure that they are able to cross the plasma membrane. Type 3 probes require additional incubation time to allow cellular esterases to remove the masking group and release the functional probe. The appropriate concentration of probe to use for a given experiment can vary depending on the specific lipid and the goals for the amount or type of protein to be extracted, and is something that researchers should be careful to optimize for their system. We recommend a 10 µM starting concentration if nothing else is known about the lipid; we provide below in the protocol some concentrations that have worked well for specific lipids. Depending on the type of lipid, and provided a very long incubation time is not required, it may be advantageous to load the probe onto cells using serum-free media, but this is something that individual researchers should test for their system. It should be mentioned that the typical alkyloxymethyl groups is cleaved more efficiently in mammalian cells than in plant cells or cells of marine species.

The coumarin photocage provides intrinsic blue fluorescence, allowing qualitative assessment of probe uptake via live-cell fluorescence microscopy. Imaging should be performed in black-walled plates to reduce background signal and photobleaching. To avoid unintended uncaging, minimal exposure to 405 nm light is recommended. Saturation is typically achieved within 2–10 minutes of treatment, though optimal probe concentration varies by compound. Uptake should be verified by comparing treated versus untreated controls. Prior studies have demonstrated successful uptake in commonly used cell lines including HeLa, HEK 293T, and Huh7^2,27,28^.

### Uncaging Efficiency

Efficient photocage removal is critical for restoring lipid bioactivity, and it is further critical that the photocage removal not begin photocrosslinking. Höglinger et al. (2017) demonstrated the orthogonality of coumarin uncaging and diazirine photocrosslinking using NMR spectroscopy before and after 405 nm irradiation^2^. To verify that probes are uncaged, thin-layer chromatography (TLC) can be used to track the generation of lipid species over time, as shown by Thomas et al. (2024)^28^. These approaches facilitate optimization of light exposure time and intensity; uncaging is typically achieved within 5 minutes using low-wattage light sources (see Supplemental Figure 4 of^16^ for a comparison of irradiation times using low-wattage light sources and high-wattage light sources).

### Crosslinking and Protein Isolation

Validation of UV-induced crosslinking at 350 to 360 nm and downstream protein enrichment is essential prior to mass spectrometry. As with uncaging, crosslinking conditions may require optimization based on the lipid probe, biological system, and light source. In-gel fluorescence, as demonstrated by Farley et al. (2024)^27^, offers a practical method to assess crosslinking efficiency. A critical consideration for preparing crosslinked lysates both for in-gel fluorescence and proteomics is the method of lysing cells. We strongly recommend not using one that relies on detergent, as this can interfere with the subsequent click reaction. We have used gentle probe sonication for best results, described below. Following irradiation and lysis, samples are reacted via click chemistry with a fluorophore-conjugated azide, resolved by SDS-PAGE, and imaged on a fluorescent scanner. This technique provides a direct measure of labeled lipid–protein conjugates and can be used to flag suboptimal or outlier samples. A small aliquot (5–10%) of each sample destined for mass spectrometry is typically sufficient for this assessment.

### Click Chemistry

These procedures require lipid-protein conjugates to be subjected to a click reaction, either to visualize the conjugates by in-gel fluorescence, or to pull down the conjugates for analysis by LC-MS/MS. The click reaction (See Fig 2B) has six components: 1) 1 mM copper sulfate 2) 1 mM sodium ascorbate 3) azide (either 20 µM fluorophore-azide or 200 µL of an azide-agarose bead slurry) 4) 100 µM TBTA 5) lysate 6) PBS. To accurately compare between samples, it is important to normalize the protein concentration in the lysate. For in-gel fluorescence, the volume of the click reaction is dictated by how much can be loaded into the well of a gel. In order to maximize the volume available, we typically run 10-well gels, often 1.5 mm thick. The total volume of our click reactions is 27 µL.

## Overview of the procedure

Below we describe three independent procedures: 1) the preparation of protein-lipid conjugates, 2) the visualization of protein-lipid conjugates via gel electrophoresis, and 3) the identification of protein interactors via proteomics analysis. Together, these enable a robust characterization of the protein interactome of select lipid species.

To begin producing protein-lipid conjugates, a cell type of interest is plated and treated with photocaged (inert) lipid probe, allowing for the probe uptake and incorporation. Upon brief irradiation with blue wavelength light (405 nm), the photolabile caging moiety is released and the probe becomes available for interactions with cellular proteins. After an incubation time to allow for probe uptake, and, for certain probes, metabolic processing, protein interactors are covalently stabilized via irradiation with near-UV light (355 nm), and cells are collected, washed, and pelleted. Discounting cell culturing time, the probe treatment and photochemistry procedure requires 2-4 hours of sample handling time.

Next, protein-lipid conjugates are visualized by in-gel fluorescence after separation by polyacrylamide gel electrophoresis (SDS-PAGE). To achieve this, cell pellets are lysed, apportioned (approximately 25 µg of input should be used for visualization, normalized across all the samples being analyzed) and subjected to a click chemistry reaction to affix a fluorophore to the terminal alkyne of the lipid probe. A standard reducing SDS-PAGE protocol has been minimally modified in order to separate the protein contents of the probe-treated samples, and the gel should be imaged on a fluorescent plate reader to visualize the fluorescent protein-lipid conjugates. The gel can then be stained with Coomassie and visualized on a standard visible-light gel imager. This procedure requires approximately 5.5-6 hours to complete.

Finally, given sufficient fluorescent signal in IGF, the remainder of each sample can be subjected to quantitative proteomic analysis to identify the protein interactors of the lipid probe. Normalized amounts of lysate are subjected to a click reaction with azide beads and then rigorously washed to remove proteins that are not covalently bound to the lipid probe. Protein-bead conjugates are then reduced, alkylated, and trypsin-digested to produce peptide mixtures which can be shipped to a proteomics facility for quantitative mass spectrometric analysis. It is highly encouraged to collaborate with the proteomics core facility on experimental design and sample preparation – specifics are beyond the scope of this protocol, though discussed in some detail below. This procedure will require approximately 10 hours across several days.

## Materials

### Reagents

- **Functional lipid probe.** Some probes are commercially available from Avanti Research, while others must be synthesized. Avanti currently offers photoactivatable and clickable (pac) probes, including pac fatty acid (Cat. #900401), pac ceramide (Cat. #900404), pac glucosylceramide (Cat. #900405), and pac phosphatidylcholine (PC; Cat. #900407). Avanti also provides select trifunctional probes (photoactivatable, clickable, and caged), including trifunctional fatty acid (Cat. #880348) and trifunctional sphingosine (Cat. #860960).
- is not an exhaustive list. We anticipate that a broader collection of published and newly developed multifunctional lipid probes will become available through *Molecular Tools Inc*., an investigator-initiated company aimed at expanding community access to these reagents, in 2026.
- Pluronic F-127 (Thermo Scientific P6867)
- HALT protease inhibitor cocktail (Thermo Scientific 87786)
- 6x Laemmli sample buffer (Thermo Scientific J61337.AD)
- TBTA (TCI T2993)
- Copper sulfate (Sigma Aldrich 61230)
- Sodium ascorbate (Sigma Aldrich A7631)
- Fluorophore-azide (Any fluorophore-azide will work; there are many commercial sources. A far red azide, such as AZDye 647 Picolyl Azide, Vector Laboratories CCT-1300, can be advantageous, as several of the ladder proteins in the BlueStain 2 protein ladder fluoresce in the far red channel).
- Picolyl Azide Agarose (Vector Laboratories, CCT-1408)
- Dithiothreitol, DTT (Sigma Aldrich D9779)
- Iodoacetamide, IAA (Sigma Aldrich I1149)
- LC-MS grade trypsin (Promega, V5071)

### Equipment

- Uncaging lamp: UV type 1 LED lamp, Nailstar model NS-02
- Photocrosslinking lamp: 37W/365 nm lamp, MelodySusie model Pro04
- Alternative lamp: Newport Lamp 1,000W Xenon-Argon lamp (with 400 and 345 nm bandpass filters)
- Probe sonicator (Many models; we used Qsonica Q55)
- 2 mL spin column (Pierce centrifuge columns: Fisher PI89896)
- C18 desalting column (BioPureSPN midi columns: Nest group HEM S18V)

### Reagent setup

- Dissolving working stocks of lipid probes (DMSO). Following manufacturer’s instructions (Avanti). 10 mM in DMSO recommended. Working stocks dissolved in DMSO can be stored at -20 °C for 6 months. It is recommended that they be aliquoted in 25-50 µL stocks. Stocks should be protected from light.
- Click reagents can be stored as concentrated stocks for up to one year at -20 °C. We maintain the following storage stocks:
- TBTA: 100 mM in DMSO
- Copper sulfate: 100 mM in water
- Sodium ascorbate: 1 M in water
- Fluorophore-azide: 1 mM in DMSO (Protect from light)
- Bead wash buffer 1 (can be stored indefinitely at room temperature):
- 100 mM Tris-HCl, pH 8
- 250 mM sodium chloride
- 5 mM EDTA
- 1 % SDS (wt/vol)
- Bead wash buffer 2 (can be stored indefinitely at room temperature)
- 100 mM Tris-HCl, pH 8
- 8 M urea
- Digestion buffer (can be stored indefinitely at room temperature)
- 100 mM Tris-HCl, pH 8
- 2 mM calcium chloride
- 10 % acetonitrile (v/v)
- 1M DTT, in water (can be stored at 4 °C for up to one year)
- 40 mM IAA, in water (can be stored at 4 °C, in the dark, for up to one year)
- Desalting Buffer 1 (80% mass-spectrometry grade ACN, 20% mass-spectrometry grade water, 0.1% TFA v/v/v) (can be stored indefinitely at room temperature)
- Desalting Buffer 2 (0.1 % TFA in mass-spectrometry grade water, v/v) (can be stored indefinitely at room temperature)
- Desalting Buffer 3 (60% mass-spectrometry grade ACN, 0.1% TFA, v/v/v) (can be stored indefinitely at room temperature)

### Biological materials

Several cell lines have been used with this procedure to generate the results described previously (see the table below) and shown in the “anticipated results” section. Below is a description of cell lines that have successfully been used with this protocol; this is not meant to be an exclusive list, as the authors have no doubt that the protocol could be applied equally well to other cell lines, with appropriate optimization of cell density and probe amount.

| Cell line | Reference | Probe |
| --- | --- | --- |
| HeLa, HEK 293T | 2 (Höglinger et al 2017) | Trifunctional fatty acid (palmitic acid, diazirine at the 11 position) |
|  |  | Trifunctional DAG |
|  |  | Trifunctional sphingosine |
| HeLa | 13 (Müller et al 2021) | PI(3,4)P<sub>2</sub> |
|  |  | PI(3,4,5)P<sub>3</sub> |
| Huh7 | 15 (Farley et al 2024) | Trifunctional fatty acid (palmitic acid, diazirine at 11 position, 5 position, or 3 position) |
| Huh7 | 21 (Farley et al 2024) | Trifunctional sphinganine |
| HeLa | 22 (Thomas et al 2025) | Trifunctional PE |
| A549 | This work (anticipated results figure) | Trifunctional fatty acid (palmitic acid, diazirine at 11 position, 5 position, or 3 position) |

### Preparing lipid-protein conjugates

#### Feeding lipids to cells

1. Seed cells the night before the experiment, with the goal of achieving around 70% confluence at the time of probe addition. The key consideration for this step is ensuring that you will recover sufficient cellular material to detect your lipid-protein conjugates, which will depend significantly on the type of cell you are using. At a minimum, you should attempt to seed enough cells that you can recover 0.3 - 0.5 mg total protein. We have found 10 cm dishes, at least one, to be an ideal size. *CRITICAL STEP: if your seeding conditions are not optimized, you will likely detect little to no signal

A. Example: A549-ACE2 cells, which are very large and not very protein-dense, required four 10 cm dishes per condition, seeded at 2 × 10^6^ cells the night before the experiment, to recover sufficient protein.
B. Example: Huh7 cells, which are much more protein dense, required one 10 cm dish per condition, seeded at 1 × 10^6^ cells the night before the experiment.
2. Prepare dilutions of probe

A. **Type 1 probes** such as fatty acids and sphingoid bases, can be diluted directly in media. While the final concentration may vary depending on your system, we have found 2 µM of sphingoid bases and 25 µM of fatty acids, diluted into complete media, to work well.
B. **Type 2 probes**, such as trifunctional PI(3,4,5)P_3_, are pre-mixed with Pluronic (10% wt/vol in DMSO) in a 1:1 ratio and then suspended in DMEM or cell supernatant to achieve the desired final concentration, usually 5 µM or 10 µM, depending on the probe and application.
3. Apply probes to cells: remove existing media from 10 cm dishes. Add 3 mL of probe-containing media to 10 cm dishes. Alternatively, remove a fraction of the cell supernatant, add to the lipid dissolved in DMSO/Pluronic and immediately add the mixture back to the 10 cm flask.
4. Incubate cells with probe, at 37 °C in 5 % CO_2_.

A. Probes that do not have a masking group (this could be a Type 1 or a Type 2 probe): incubate for 10-30 minutes, then remove the probe solution and replace with complete media.
B. **Type 3 probes**, such as trifunctional PI(3,4,5)P3 require more time for cellular esterases to remove the masking group: incubate overnight, then remove the probe solution and replace with complete media.

*TROUBLESHOOTING*

This protocol must be adjusted for the particularities of the cell type being analyzed. For example, high-adherence plating conditions (e.g. poly-L-lysine coated dishes) should be used for poorly adherent cell lines (e.g. HEK293T); similarly, the media used to dilute the lipid probe should be matched to the cell type (e.g. DMEM versus RPMI or Astrocyte Medium).

#### Photochemistry and cell harvesting

5. Uncage: if the probe carries a caging group, expose dishes, uncovered, to 400 nm light for five minutes. Potential sources for 400 nm light are discussed above (Equipment).
6. Photocrosslink: expose dishes, uncovered, to 350 nm light for five minutes. Potential sources for 350 nm light are discussed above (Equipment).
7. Immediately after photocrosslinking, place dishes on ice. *CRITICAL STEP* This and all subsequent steps should be performed on ice and/or at 4 °C.
8. Decant media from dish
9. Wash dishes, gently, with pre-chilled PBS
10. Scrape cells into 2 mL with a cell lifter and transfer to a 15 mL conical tube. Rinse the plate with 2 mL PBS to collect any remaining cellular material and add it to the 15 mL tube.
11. Spin down cells into a pellet: centrifuge at 1,000 x g for 5 minutes
12. Decant supernatant
13. Resuspend cell pellet in PBS. The volume of PBS used for this resuspension will dictate the maximum concentration of the protein solution you are able to use in subsequent steps. 250-1,000 µL is recommended (500 µL is a good place to start if you have not yet optimized this step).

*TROUBLESHOOTING*

14. Transfer the resuspended pellet to a 1.5 mL microcentrifuge tube.
15. *PAUSE POINT* at this point, the resuspended pellet can be stored at 20 °C for up to six months

### Visualizing protein-lipid conjugates by in-gel fluorescence

#### Lysis

16. Thaw cell pellet, on ice. Once cell pellet is thawed, add a protease inhibitor cocktail, such as the HALT protease inhibitor cocktail referenced in the MATERIALS.
17. Still on ice, sonicate each sample on the lowest possible power setting, in 3 10-second bursts, allowing the sample to cool in between bursts. The liquid should be perfectly clear at the end of successful lysis.

*CRITICAL STEP protein yield will be significantly damaged if the sample is over-sonicated, or if protease inhibitor cocktail is not used.

*TROUBLESHOOTING*

18. Determine the concentration of each sample. This can be done with any protein concentration assay (BCA, Bradford); we do not recommend attempting to determine protein concentration by absorbance-based methods (eg, Nanodrop).

#### Click reaction

19. Prepare dilutions of each sample lysate in PBS, normalizing all samples to contain the same amount of protein (see example table below). The total volume for each sample of lysate should be 25 µL.

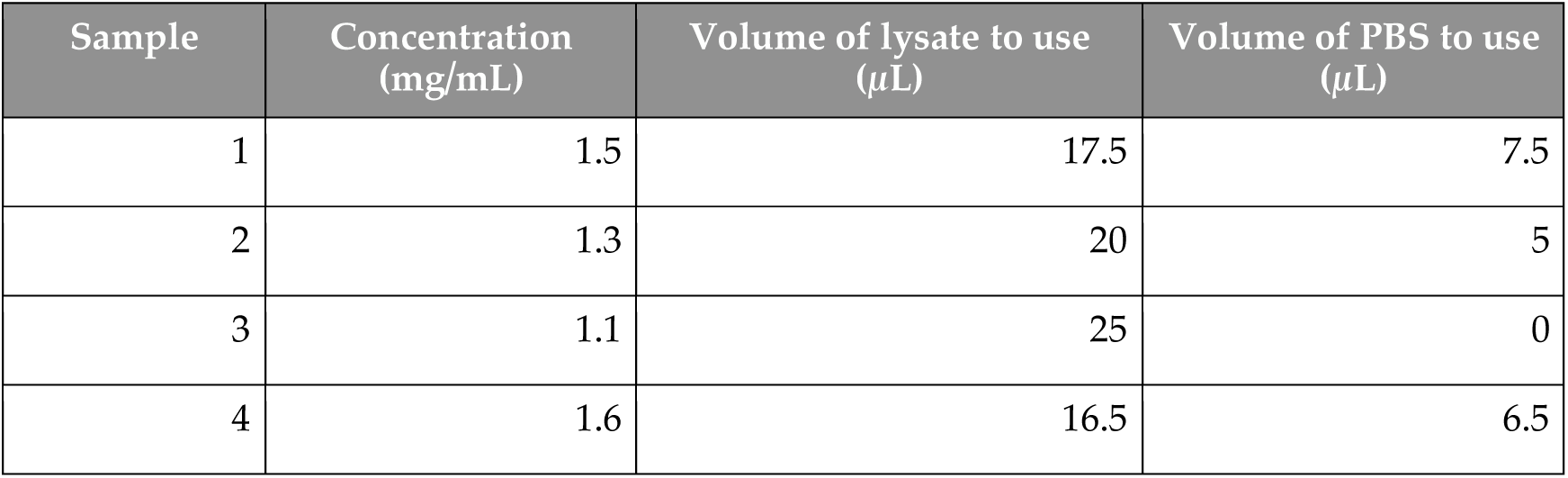

| Sample | Concentration (mg/mL) | Volume of lysate to use (μL) | Volume of PBS to use (μL) |
| --- | --- | --- | --- |
| 1 | 1.5 | 17.5 | 7.5 |
| 2 | 1.3 | 20 | 5 |
| 3 | 1.1 | 25 | 0 |
| 4 | 1.6 | 16.5 | 6.5 |

20. Prepare the master mix of click reagents (TBTA, copper sulfate, sodium ascorbate, and fluorophore-azide). We recommend preparing working stocks of each reagent so that equal volumes of each can be added to the master mix (see example table below). We have not found that the order of addition of the reagents to the master mix makes any difference whatsoever. First adding your reducing agent (sodium ascorbate) to copper sulfate, however, can be a way reassure yourself that the reduction is happening properly, as an instantaneous color change from blue copper II to yellowish copper I is visible if the reagents are concentrated enough.

| Reagent | Stock concentration | Goal concentration | Volume (1 sample) | Number of Samples | Volume for master mix |
| --- | --- | --- | --- | --- | --- |
| TBTA | 5.4 mM | 100 μM | 0.5 μL | 10 | 5 μL |
| CuSO<sub>4</sub> | 54 mM | 1 mM | 0.5 μL | 10 | 5 μL |
| Sodium ascorbate | 54 mM | 1 mM | 0.5 μL | 10 | 5 μL |
| Fluorophore-azide | 54 mM | 1 mM | 0.5 μL | 10 | 5 μL |

21. Add 2 µL of click master mix to 25 µL of diluted lysate. Incubate in the dark at room temperature for 1 hour, with gentle shaking/rocking

#### SDS-PAGE analysis

22. Quench the click reaction by adding 5.4 µL of a 6x stock of Laemmli buffer
23. Incubate samples at 42 °C for 30 minutes (Note: do **not** boil samples; boiling samples will cause membrane proteins to crash out and aggregate. Solutions of membrane proteins should never be heated above 60 °C.)
24. Run SDS-PAGE gel, selecting an acrylamide concentration appropriate to the protein(s) of interest.
25. *CRITICAL STEP: Visualize the fluorescence using a fluorescent imager; a wide range of imagers are appropriate, depending on the excitation/emission requirements of the fluorophore selected. A far-red fluorophore has the advantage of being able to be visualized alongside the BlueStain2 protein ladder. **Note**: When performing in-gel fluorescence, use a compatible protein ladder (e.g., Blue Stain) and choose a fluorophore–azide whose emission does not overlap with the ladder dye to avoid signal interference.
26. If desired, stain the gel with Coomassie (or other desired protein stain) to visualize total protein. **Note**: do **not** stain prior to visualizing fluorescence.

### Identifying lipid-binding proteins by tandem mass spectrometry

#### Click reaction

27. This part of the procedure starts from the lysate prepared and analyzed in steps 16-18. For each sample, transfer 200 µL of picolyl-azide bead slurry to a 1.5 mL microcentrifuge tube. (\***CRITICAL STEP**: before the transfer, ensure the bead slurry is well-homogenized by vortexing. If the bead slurry is not well-homogenized, different amounts of beads will end up in each sample tube, introducing artificial bias in the amount of protein that is ultimately pulled down).
28. Wash beads by centrifuging (1,000 x g for 2 minutes), carefully removing supernatant, adding 500 µL water, centrifuging again, and carefully removing supernatant. Since beads are initially a 50/50 v/v slurry, about 100 µL of beads should remain.
29. Prepare master mix of click reagents (TBTA, copper sulfate, and sodium ascorbate). Recommended working stocks and dilutions are described in the table below

| Reagent | Stock concentration | Goal concentration | Volume (1 sample) | Number of Samples | Volume for master mix |
| --- | --- | --- | --- | --- | --- |
| TBTA | 100 mM | 100 μM | 1 μL | 10 | 10 μL |
| CuSO<sub>4</sub> | 100 mM | 1 mM | 10 μL | 10 | 100 μL |
| Sodium ascorbate | 1 M | 1 mM | 1 μL | 10 | 10 μL |

30. To the 100 µL of pre-washed azide beads, add 500 µL diluted lysate, 388 µL of PBS, and 12 µL of click master mix.
31. Rock tubes gently at room temperature for 1 hour. An end-over-end turning situation is ideal; it is critical that the beads not be allowed to settle, and instead be consistently moved through the lysate. After this step, your tubes should contain lipid-bound proteins covalently linked to the azide beads, and all the other proteins from the lysate should be in solution.

#### Sample preparation for proteomics

32. In order to confidently characterize proteins as lipid binding, it is essential to stringently wash the azide beads to remove proteins that are not covalently linked. Centrifuge beads (1,000 x g for 2 minutes, for this and all subsequent steps). Wash beads once with PBS, and then resuspend in 500 µL PBS. Transfer the slurry to a 2 mL spin column. Use another 500 µL PBS to rinse the centrifuge tube and transfer to the 2 mL spin column, to ensure that all beads were transferred.
33. Wash five times with 2 mL bead wash buffer 1
34. Wash ten times with 2 mL bead wash buffer 2
35. Remove the beads from the column in 500 µL PBS and transfer to a fresh microcentrifuge tube. Rinse column with PBS to ensure all beads have been transferred. Spin down beads and carefully remove supernatant.

*TROUBLESHOOTING*

36. **Reduce** proteins with dithiothritol (DTT), by resuspending beads in 1 mL digestion buffer and adding 10 µL of 1 M DTT in water. Incubate beads at 42 °C for 30 minutes.
37. Spin down beads and carefully remove supernatant.
38. **Alkylate** proteins with iodoacetamide (IAA), by resuspending them in 1 mL of 40 mM IAA. Incubate at room temperature in the dark for 30 minutes.
39. Spin down beads and carefully remove supernatant.
40. **Digest** beads by adding 50 µL of digestion buffer + 1 µL LC-MS-grade trypsin (resuspended following manufacturer’s instructions). Shake at 37 °C overnight.
41. Activate a C18 desalting column with 400 µL methanol, centrifuge for 2 minutes at 1,000 x g (for this and all subsequent steps)
42. Wash the column 3 x 300 µL with Desalting Buffer 1
43. Wash the column 3 x 300 µL with Desalting Buffer 2
44. Load the sample (∼150 µL beads + supernatant). Spin down once.
45. Wash column 3 x 300 µL with Desalting Buffer 2
46. Move column to fresh microcentrifuge tube. Elute 2 x 200 µL with Desalting Buffer 3.
47. Use a speedvac (preferably cooled) to dry samples. Dried peptides can be stored at -20 °C and shipped on dry ice to a proteomics facility.

(*PAUSE POINT: dried peptides can be stored at -20 °C for up to six months)

### Proteomics

A comprehensive overview of mass spectrometry (MS)-based proteomics is beyond the scope of this protocol, as the field is highly specialized and continues to evolve rapidly. Numerous excellent resources provide in-depth coverage of foundational and emerging proteomics strategies. Here, we outline several established quantitative proteomics approaches that have been successfully applied to analyze protein–lipid interactions (Table 4). These methods are not uniquely tailored to lipid interactome studies but instead leverage broadly applicable MS-based workflows originally developed for general proteomic analyses.

**Table 4.** Overview of strengths and weaknesses of common quantitative mass spectrometry techniques. Abbreviations: PSM = peptide spectral matching; SILAC = Stable Isotope Labeling by Amino acids in Cell culture; iTRAQ = Isobaric tags for relative and absolute quantitation; TMT = Tandem Mass Tagging; DIA LFQ = Data Independent Acquisition Label Free Quantitation.

| Method | # Samples per cassette | Major strengths | Major weaknesses | Methodology references |
| --- | --- | --- | --- | --- |
| PSM | Any | Highly affordable, applicable on many MS instruments.<br>Widely established.<br>New computational methods are improving the interpretation of this data. | Weak quantitation.<br>Requires longer instrument run times.<br>Field is leaving this method behind. | 33,34 |
| SILAC | 2 or 3 | Effective ratiometric quantification, applicable on many MS instruments<br>Widely established.<br>Lower risk of technical variation between samples. | Smaller theoretical maximum for number of protein IDs.<br>Relatively expensive.<br>Reliant on cells/tissues with high turnover rate.<br>Field is leaving this method behind. | 35,36 |
| iTRAQ | 4 or 8 | Robust quantitation.<br>Requires moderate resolution MS instrument.<br>Allows for significant multiplexing. | Expensive for number of samples/cassette.<br>Offers little benefit over higher multiplexed methods such as TMT. | 37,38 |
| TMT | 6, 16, 35 | Extremely robust ratiometric quantitation.<br>Widely used.<br>Affordable on a per-sample basis.<br>High multiplexing reduces instrument time. | Relatively high ratiometric compression due to bleedover/contamination of adjacent channels.<br>Relatively higher cost.<br>Requires high-resolution MS instrument | 37,38 |
| DIA LFQ | Any | High sample throughput.<br>Robust ratiometric quantification.<br>Comprehensive data acquisition (reduced missing data based on peptide abundance). | Methodology still under development, requires high-resolution MS instrument, each sample run separately resulting in longer instrument running time | 39,40 |

Several quantitative strategies have been employed to map lipid-associated protein networks. Label-free approaches such as **spectral counting** offer a relatively simple and accessible means to estimate protein abundance based on the number of identified peptide spectra per protein. This method has been used to study the interactomes of bifunctional fatty acids, bifunctional sphingosine, and trifunctional sphingosine^27,29,30^.

However, spectral counting can be limited in dynamic range and may lack the sensitivity required to robustly quantify low-abundance interactors.

More quantitative and precise techniques include **stable isotope labeling by amino acids in cell culture (SILAC)**, which enables direct comparison of differentially treated samples by incorporating heavy isotopes (e.g., ¹³C, ¹⁵N) into proteins during cellular metabolism. SILAC has been effectively used to investigate protein–lipid interactions involving bifunctional endocannabinoids and sterols^5,31^, offering improved reproducibility and reduced technical variability compared to label-free methods.

**Isobaric tagging techniques**, such as **tandem mass tag (TMT)** labeling, have become increasingly popular due to their high multiplexing capacity. TMT allows simultaneous analysis of up to 32 samples in a single experiment, making it particularly well-suited for complex study designs such as time-course or dose-response experiments. This strategy has been applied to quantify interactomes involving trifunctional sphinganine and phosphatidylethanolamine^27,32^, demonstrating its utility in high-throughput lipid–protein interaction studies.

Recently, **data-independent acquisition (DIA)** has emerged as a powerful alternative to traditional data-dependent acquisition (DDA), offering improved reproducibility and quantitative accuracy across complex samples. By systematically fragmenting all precursors within defined mass windows, DIA enables comprehensive and unbiased proteome coverage. Its adoption in the field of interactomics, including studies of membrane-associated and lipid-binding proteins, is growing rapidly and is expected to further enhance the depth and consistency of lipid-interactome mapping in the near future.

In conclusion, no single proteomics strategy is universally optimal for analyzing protein–lipid interactions. Each method presents distinct advantages and trade-offs in terms of sensitivity, throughput, quantitative accuracy, cost, and required expertise. The choice of approach should be guided by the specific biological question, sample type, experimental complexity, and the analytical platforms available within a given laboratory. As the field of proteomics continues to evolve, driven by technological innovations in instrumentation, labeling chemistry, and computational analysis, researchers are encouraged to stay abreast of recent developments and validate key findings using orthogonal approaches whenever possible.

### Timing

Step 1, cell seeding: 1 hour (see Figure 3)

**Figure 3:**
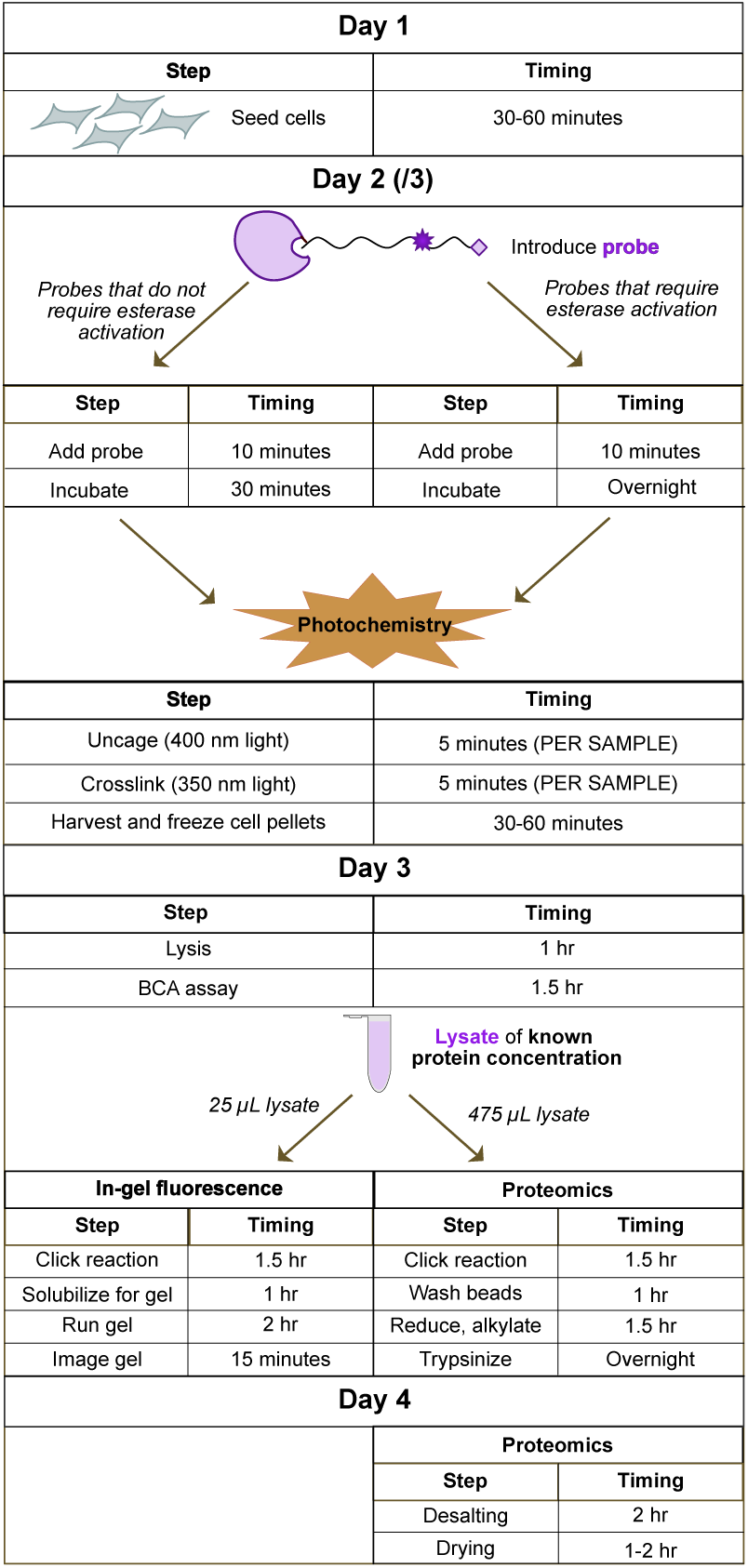
Summary of timing for the workflow in Protocols 1, 2, and 3. While Pause Points are indicated in the protocols above, we present here what a theoretical experiment would look like if it were to be completed continuously. We furthermore indicate what it would look like to simultaneously run a gel and proteomics on the same sample.

Step 2-4, probe application and sample processing: 2-4 hours. Note: for probes that require esterase activation, an overnight incubation with probe is required, spreading the work for step 4 across 2 days.

Step 5: 5 minutes per 10 cm dish

Step 6-7: 5 minutes per 10 cm dish (multiplexing using multiple light sources can reduce the time required for steps 5 and 6, if large numbers of dishes are being processed)

Steps 8-15, cell harvesting: again, depends on the number of plates, usually 30-60 minutes.

Steps 16-17, lysis: 1 hour (including thawing)

Step 18, concentration determination: 1.5 hours

Steps 19-21, click reaction for in-gel fluorescence: 1.5 hours

Steps 22-26, SDS-PAGE running and visualization: 3 hours

Steps 27-31, click reaction for proteomics: 1.5 hours

Steps 32-35, bead washing: 1 hour

Steps 36-40, reduction, alkylation, digestion: 2 hours [**ends with overnight step**] Steps 41-47: 4 hours

### Troubleshooting

Most problems that occur within the protocol cannot be observed until the very end, with the demoralizing result of a blank gel or uninterpretable proteomics results, and it can be difficult to trace back where the problem arose. Two pieces of general good practice will help to reduce at least some confusion about the source of errors. #1: always take along appropriate controls. At a minimum, you should include samples that were photocrosslinked to produce protein-lipid conjugates, and samples that were NOT exposed to 350 nm light for photocrosslinking. This will allow you to assess background phenomena such as leaky (ambient) photocrosslinking and insufficient bead washing. For a pure assessment of true background you can also include samples to which no lipid probe was added. #2: always run a gel in parallel, of any samples that you are analyzing by mass spectrometry. As depicted in the “Timing” figure, it is straightforward to save 25 µL of lysate to run on a gel while the rest of the material goes on for mass spectrometry analysis. This will allow a quick visual assessment of whether sufficient protein was obtained, and whether sufficient difference between (+) UV and (-) UV samples was observed. Examples of problems that you may experience are described below:

| Step | Problem | Possible reason | Solution |
| --- | --- | --- | --- |
| 2-4 | Fluorescent bands on gel are indistinct or absent | Probe was not loaded properly and did not enter cells | If probes are caged, it is possible to visually assess whether probe entered cells by live-cell microscopy monitoring the coumarin channel. If probe did not enter cells, increase one or all of 1) concentration of probe 2) volume of probe solution applied to cells 3) amount of time probe sits on cells |
| 1 | Fluorescent bands on gel indistinct or absent | Cell density is not high enough - not enough protein is present to detect signal | If probes are entering cells correctly (as assessed above), you may need to increase the protein concentration of the sample you run the click reaction on. Try using more dishes of cells and/or lysing in a smaller volume |
| 16-17 | Fluorescent bands are weak and/or concentrated at the stacking gel/running gel interface | Lysis was incomplete | Include more bursts of sonication if there are visible precipitates in solution, or include a complementary method of lysis (eg by passing cell suspension through a low-gauge needle) |
| 22 | Fluorescent bands are weak and/or concentrated at the stacking gel/running gel interface | Proteins were not well-solubilized | Increase amount of time post-click lysates are allowed to incubate at 42 °C, or increase temperature to (but not above!) 60 °C. |
| 31-35 | Dynamic range between (+) UV and (-) UV samples in proteomics analysis is very low | Insufficient washing of beads | If signal in proteomics is low, but the bands in the gel show a robust difference between bands that fluoresce in the (+) UV and (-) UV samples, there is likely a high degree of background contamination. You can increase the number of times you wash the beads with buffers 1 and 2. |
| 27-31 | Dynamic range between (+) UV and (-) UV samples in proteomics analysis is very low | Insufficient solubilization or click reaction | It is also possible that the click reaction with the beads failed due to insufficient contact between azide groups and exposed alkyne. This can occur if proteins are not well-solubilized (eg, if there are visible precipitates) or if there was insufficient rocking during the click reaction |

## Anticipated results

While the goals of an experiment profiling lipid-protein interactions may vary, the basic requirements are 1) detectable signal (either fluorescence in a gel or intensity of relevant fragment ions in mass spectrometry) and 2) differences in intensity between samples irradiated with 350 nm light for photocrosslinking, and samples not exposed to crosslinking radiation. In Figure 4, we present examples of experiments throughout the optimization process to produce these two outcomes. Low initial fluorescent signal in gel samples (Figure 4A) indicated that fundamental issues were present in the experimental design. By increasing the amount of time cells were incubated with probe, and improving lysis protocols, we were able to observe detectable signal (Fig 4B). We tested multiple seeding conditions for our cell lines, and selected one that produced the most robust fluorescent signal. We finally arrived at optimal lysis conditions and guidelines for protein concentration that we report in the above protocol, which reproducibly produce gels with robust signal intensity, and dramatic differences in intensity between +/- UV conditions, as displayed in Fig 4C. Fig 4C has been published, in reference^16^.

**Figure 4:**
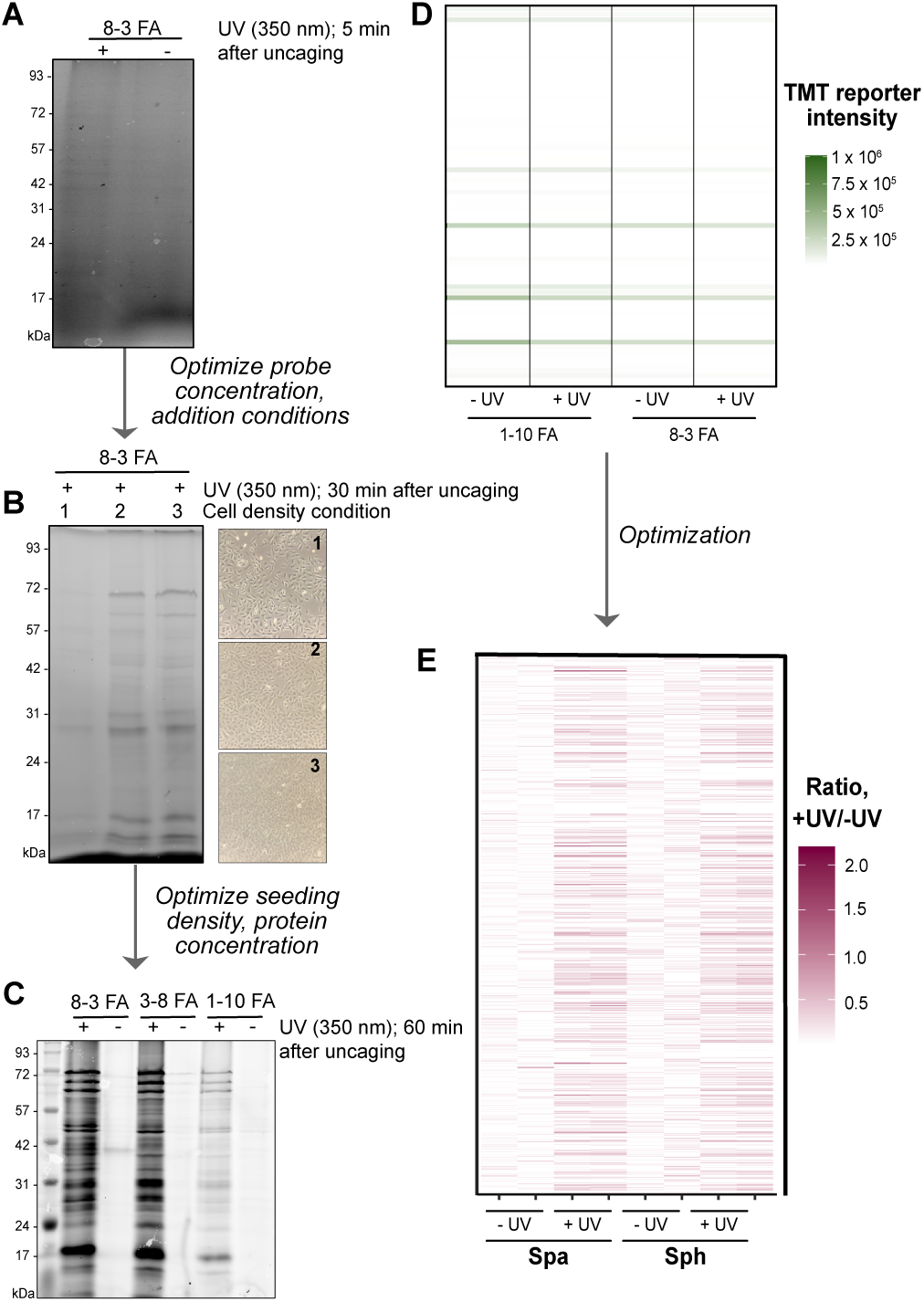
Anticipated results. **(A)** A549 cells incubated with a trifunctional fatty acid probe and either irradiated with 350 nm light or left in the dark. 25 μM 8-3 FA was incubated for only 5 minutes on cells, in a volume of 1 mL, and cells were lysed by mechanical disruption (passing through an 18-gauge needle). **(B)** A549 cells incubated with a trifunctional fatty acid probe and irradiated with 350 nm light. Three densities of A549 cells were used, corresponding to image 1, 2 or 3. Images were taken with a cell phone camera through a simple light microscope at 20x magnification. 25 μM 8-3 FA was incubated for 30 minutes on cells, in a volume of 3 mL, and cells were lysed by sonication, as described in Protocol 1 above. **(C)** A549 cells were incubated with three different trifunctional fatty acid probes (all based on palmitate, with the diazirine at different locations on the acyl chain) and either irradiated with 350 nm light or left in the dark. Gel was obtained following Protocols 1 and 2 above. **(D)** Proteomics of Huh7 cells treated with two trifunctional fatty acid probes (both based on palmitate, with the diazirine at different locations on the acyl chain) and either irradiated with 350 nm light or left in the dark, harvested following the protocols that produced the gel in (A). **(E)** Proteomics of Huh7 cells treated with trifunctional sphinganine (Spa) or trifunctional sphingosine (Sph), harvested and processed following Protocol 1 and Protocol 3. Fig 3C is reproduced from reference^16^; Fig 3e is reproduced from reference^27^.

Similar optimization is required for proteomics experiments. A poor proteomics experiment, as shown in Fig 3d, indicates little to no difference in signal between +/- UV conditions, indicating high contamination of background proteins, little enrichment of lipid-binding proteins, or both. Similar optimization of probe addition, cell density, and lysis as described above, as well as improved bead-washing protocols, resulted in a more dramatic difference between +/- UV conditions for multiple lipid probes, as shown in Fig 4E. Fig 4E has been published, in reference^27^.

## Hit validation

As with any interactome study, lipid–protein interactome datasets can yield a large number of candidate hits. Robust statistical analysis is essential to prioritize high-confidence interactors, including approaches that account for peptide coverage, reproducibility across replicates, and appropriate significance metrics (e.g., adjusted p-values). Equally important, prioritized hits should be validated using independent, orthogonal methods. Validation can include direct biochemical confirmation of lipid association for individual candidates. In our own studies, we used targeted pull-down followed by western blot to confirm PITPNB as an interactor identified with a functionalized fatty-acid probe^16^, and similarly validated VDAC2 as a ceramide-interacting protein^41^. Additional orthogonal approaches may include recombinant protein binding assays, co-fractionation or co-localization studies, and competition experiments using the corresponding unlabeled lipid to assess specificity.

Beyond binding, functional validation helps establish biological relevance. This can be done by genetic perturbation (CRISPR–Cas9 knockout, CRISPRi, or siRNA knockdown) to test whether loss of the candidate alters the phenotype or pathway linked to the lipid of interest. Conversely, overexpression of the candidate protein can be used to assess effects on activity, lipid-dependent signaling, or subcellular localization.

Together, these steps help triage large hit lists and distinguish true, functional lipid–protein interactions from background.

## Acknowledgments

This work is supported by NIH NIAID grant R01 AI141549 (F. G. T. and C. S.), and NIH NIGMS R01 GM127631 and R35 GM158174 (C. S.), an Allen Distinguished Investigator Award provided by a Paul G. Allen Frontiers Group advised grant of Allen Family Philanthropies (C. S.).

